# Investigating the Zoonotic Origin of the Marburg Virus Outbreak in Guinea in 2021

**DOI:** 10.1101/2022.11.03.514981

**Authors:** Marat Makenov, Sanaba Boumbaly, Faya Raphael Tolno, Nouminy Sacko, Leno Tamba N’Fatoma, Oumar Mansare, Bonaventure Kolie, Olga Stukolova, Evgeny Morozkin, Ivan Kholodilov, Olga Zhurenkova, Marina Fyodorova, Vasily Akimkin, Anna Popova, Namoudou Conde, Mamadou Yero Boiro, Lyudmila Karan

## Abstract

In 2021, a patient died from Marburg virus (MARV) disease in Guinea, which was the first confirmed case in West Africa. The source of the outbreak has not been identified. It was revealed only that the patient had not traveled anywhere before the illness. Prior to this outbreak, MARV had been found in bats in the neighboring Sierra Leone. In Guinea, this virus has never been found before. Therefore, the question of the source of infection arose: was it an autochthonous case with spillover from a local population of bats or an imported case with spillover from fruit bats foraging/migrating from Sierra Leone?

In this paper, we aimed to conduct a study of *Rousettus aegyptiacus* in Guinea and determine the source of the most likely infection of a patient who died from Marburg virus disease in 2021 in Guinea.

We caught bats at 32 sites in Guéckédou prefecture, including seven caves and 25 locations on the flight path. A total of 501 fruit bats (Pteropodidae) were captured, including 66 *R. aegyptiacus*. The subsequent screening showed three PCR-positive MARV bats. We have found and described two caves in Guéckédou prefecture where MARV-positive *R. aegyptiacus* roost. Sequencing has shown that at least two different MARV genetic variants circulate in *R. aegyptiacus* in Guinea: an Angola-like strain and MARV strains of major marburgvirus lineages.

## Introduction

In August 2021, in Guinea, in Temessadou M’bokét village (Guéckédou prefecture), a patient presented to a local health care point with signs and symptoms of hemorrhagic fever. The patient died the next day in the village. This case was investigated and shown to be Marburg virus disease (MARD) [1]. Sequencing of an isolate from the Guinean patient showed that this outbreak was caused by the Angola-like

Marburg virus (MARV) [1]. Previously, in 2017-2018, MARV was detected in the Egyptian rousette bat (ERB; *Rousettus aegyptiacus*) in the neighboring Sierra Leone [2]. This was the first detection of MARV in West Africa. Phylogenetic analysis revealed that at least two genetic variants of MARV circulate in bats in Sierra Leone: Angola-like MARV and MARV, which are genetically similar to variants from Uganda, Gabon, and the Democratic Republic of Congo (DRC) [2].

MARV belongs to the *Filoviridae* family and causes a severe hemorrhagic infection in humans with a high mortality rate [3]. The Angola-like variant became known after the largest outbreak of MARV, which occurred in 2005 in Angola, with 252 cases and 227 deaths [4]. The reservoir of MARV is ERB [5]. These fruit bats live in colonies with a large number of individuals and form roosts in caves [6,7]. In addition, MARV was detected in the green monkey *Chlorocebus* sp. [8,9], and in the cave-dwelling insectivorous bats *Miniopterus inflatus* and *Rhinolophus eloquens* [10].

Thirteen MARD outbreaks had been described up to 2019 [11,12]. All of them (with the exception of two outbreaks of laboratory infection in Germany and Russia) were of African origin. Epidemiological investigations of these outbreaks showed that seven MARD outbreaks (from 13) emerged after the index patient visited a cave or mine, and their location was precisely determined [11]. To date, the list of such caves includes Sinoia cave in Zimbabwe, Kitum cave in Kenya, Python cave and Kitaka cave in Uganda [5,13–15]. The presence of large colonies of bats in these caves has been confirmed. In some caves, MARV spillover events have occurred repeatedly; in particular, two outbreaks emerged after visiting Kitum cave in 1980 and 1987 (Kenya) [13,16], and two outbreaks emerged after visiting the Python cave in 2008 (Uganda) [14]. However, for some MARD outbreaks, including the most numerous outbreak in Angola in 2004-2005, the origin of human infection has not been identified [4].

In the case of the patient from Temessadou M’bokét, it was not possible to know for sure whether he visited such caves or had contact with bats before the disease. Long-distance flights are known for ERB: their maximum foraging distance varies between 11-32 km, and the confirmed maximum relocation distance was 500 km [17–19]. The Koema Cave (Sierra Leone) where MARV has been found is 56 km away from Temessadou M’bokét village. This distance fits well with the movement ecology of ERB, and it allows us to assume that the MARV could have been introduced in Guinea with MARV-positive ERB from Sierra Leone. In contrast, it is possible that MARV had not been introduced from Sierra Leone and that there are caves in Guinea that are also inhabited by MARV-positive ERB. Finding ERB roosting caves, as well as determining their status according to MARV, is the most important task to prevent new MARD outbreaks.

In Guinea, the largest Ebola virus disease outbreak emerged in 2013, which increased interest in the search for filoviruses in animals. Saéz et al. [20] attempted to find the source of Ebola virus spillover from a natural source [20]. The authors began their study in Guéckédou prefecture in the village of Meliandou, where the index patient lived and died. They caught and screened 169 bats of 13 species and obtained negative results [20]. In addition, screening of bats for filoviruses was carried out in Guinea [21] after the discovery of the Bombali virus (*Bombali ebolavirus*) in neighboring Sierra Leone [22]. The MARD outbreak in Guinea in 2021 brought scientists back to the search for filoviruses in this country; only this time the search involved not only the genus *Ebolavirus* but also the genus *Marburgvirus*. A team of WHO specialists arrived in Temessadou M’bokét in August-September 2021 for an epidemiological investigation of the case and searched for the MARV reservoir [23]. The team conducted bat trapping in the immediate vicinity of the village of Temessadou M’bokét, as well as in the vicinity of neighboring villages (Baladou Pebal and Koundou) [23]. However, publications on the detection of MARV-positive bats in Guinea have not followed.

In this paper, we aimed to conduct a study of *R. aegyptiacus* in Guéckédou prefecture and determine the source of the most likely infection of a patient who died from MARD in 2021 in Guinea.

## Methods

### Bat capture and processing

We captured animals in June-July 2022. A two-step strategy was chosen to search for the ERB roosting caves. First, bats were captured on flight paths using mist nets. Second, we searched for caves and caught bats of the inhabitants of these caves by setting nets at the entrances and in close proximity to the caves. The search for caves was carried out based on the data of bat captures on flight paths: if ERB were present in the captures, then we came to nearby villages and asked local residents whether there were caves in the vicinity.

Bat species were identified based on morphology [37]. We divided all caught ERB into two age groups: juveniles (under 6 months of age) with forearm lengths less than 90 mm and adults with forearm lengths greater than 89 mm. Standard methods were followed for the safe handling and sampling of mammals that are potentially infected with infectious pathogens [38]. Captured bats were placed in breathable cotton bags, which were moisturized to prevent bat dehydration and transported to the field laboratory. Then, the bats were euthanized according to standard protocols. Blood from the euthanized bats was collected through cardiac puncture into sterile tubes with 0.5 M EDTA. Blood samples were centrifuged to separate plasma and cells. Urine was collected antemortem in sterile Petri dishes or, postmortem, with a sterile syringe from the bladder. During necropsy, samples of skin, sections of the brain, lymph nodes, salivary glands, liver, spleen, kidney, lung, and intestines were obtained and stored in RNA tissue protect buffer (Qiagen, Hilden, Germany) at – 20°C.

Pools of internal organs (liver, spleen) and lymph nodes of bats were homogenized with TissueLyser LT (Qiagen, Hilden, Germany) in 0.5 ml of 0.15 M NaCl solution immediately after necropsy. Total RNA was extracted from 50 μl of 10% suspension using an RNeasy Plus kit (Qiagen, Hilden, Germany). For PCR-positive samples, skin homogenates, ectoparasites, oral swabs, blood plasma, urine, and nonpooled organ tissues (brain, liver, spleen, lung, kidney, lymph nodes, and salivary glands) were additionally studied using the same preparation and RNA extraction methods.

### Viral detection and sequencing

Detection of filovirus RNA was performed using an Amplisens FiloA-Fl commercial kit (Central Research Institute of Epidemiology, Moscow, Russia) designed for the detection of *Zaire ebolavirus, Sudan ebolavirus*, and Marburg virus. Positive samples were reverse transcribed using a Reverta-L RT kit (Central Research Institute of Epidemiology, Moscow, Russia) according to the manufacturer’s instructions.

Four gene fragments of MARV were Sanger sequenced using the primers shown in Table 1. One PCR-positive sample with a low RNA concentration was sequenced only with the primers used in the qRT□PCR screening. Possible contamination was ruled out by comparison of the obtained fragment sequence with the sequence of the recombinant positive control used in the PCR kit. The purified PCR products were sequenced bidirectionally using a BigDye Terminator v1.1 Cycle Sequencing kit (Thermo Fisher Scientific, Austin, TX, USA) on an Applied Biosystems 3500xL Genetic Analyzer (Applied Biosystems, Foster City, CA, USA). The obtained nucleotide sequences were aligned, compared, and analyzed using MEGA X [39].

**Table 1.**
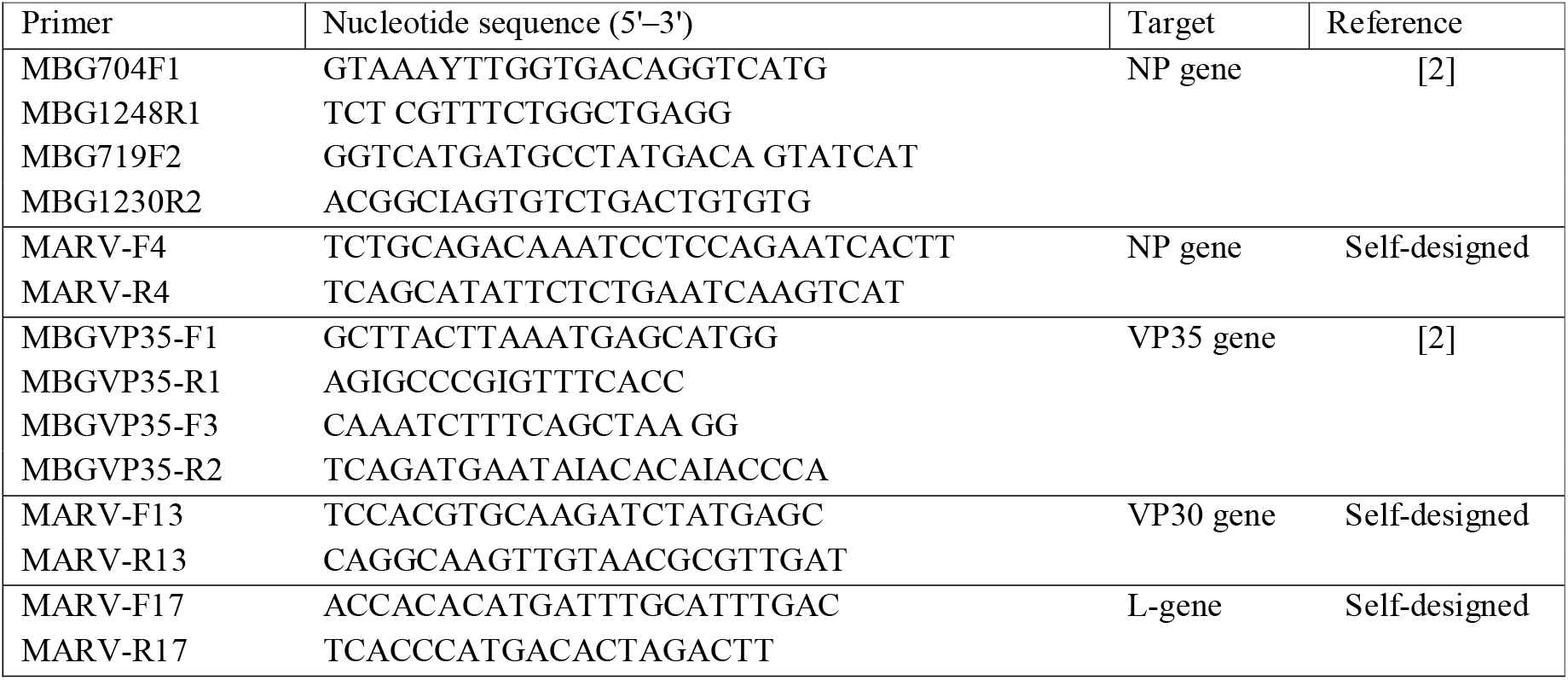
Primers used in sequencing.

### Ethics Statement

The procedures used in this study adhered to the tenets of the Declaration of Helsinki. Approval was obtained from the National ethics committee for Health Research of Guinea (document number 061/CNERS/15), and from Research Institute of Applied Biology of Guinea (document number 14/DG/IPG/K/2019).

## Results

### Searching for *Rousettus aegyptiacus* daily roost caves

During this study, we caught bats at 32 sites in Guéckédou prefecture, including seven caves and 25 locations on the flight path. A total of 501 fruit bats (Pteropodidae) were captured, including *Lissonycteris angolensis* (224 bats), *Myonycteris leptodon* (110 bats), *R. aegyptiacus* (66 bats), *Epomops buettikoferi* (82 bats), *Hypsignathus monstrosus* (13 bats), *Eidolon helvum* (4 bats), and *Micropteropus pusillus* (2 bats). We started our investigation from the Temessadou M’bokét cave, which is closest to the Temessadou M’bokét village.

#### Temessadou M’bokét cave

(N 8.61706°, W 10.36345°) is located 600 m from Temessadou M’bokét village, where an MARV-positive patient died in 2021. It is a shallow talus cave that was formed among large boulders. Examination of the cave and captures at its entrances revealed that it was inhabited only by *L. angolensis*, *Hipposideros caffer/ruber*, and *Hipposideros beatus*. ERB were not found in the cave. It is known that *R. aegyptiacus* prefer deep and dark caves for roosting in the daytime and do not inhabit talus caves [6], so we assume that ERB were absent not only at the time of our study, but in general, as this cave was not attractive for ERB. Therefore, it is unlikely that the MARV-positive patient became infected while visiting this cave.

After investigation of this first cave, we switched to a two-step algorithm in which the first bats were trapped in flight paths, and after capturing the ERB, the surroundings were investigated for the presence of caves. Using this strategy, ERB were trapped in flight paths in eight different locations, and two caves were found in which there were ERB roosts at the time of the study.

#### Legon Tyo cave

(N 8.64561°, W 10.37876°) is located in Koundou subprefecture, close to Tongolo village (~600 m), on the northern slope of Legon Tyo Mountain. Near the entrance to the cave, the mountainside is covered with small plantations of rice, pepper, cassava and other plants cultivated by locals. There are several talus caves on the slope in front of the entrance to the main cave. *M. leptodon* and *L. angolensis* roosted in these small cavities between large boulders. The entrance to the main cave is located 50 meters above these talus caves on an almost sheer slope and is not accessible to people. We caught 19 ERB near the entrance to the cave, as well as in the flight paths in the vicinity of the cave (Table 2). Temessadou M’bokét village, where the 2021 MARD outbreak emerged, is 4.5 km away from this cave. An important detail is that the cave is located on the lands of the farmers of Tongolo village, and the passage of outsiders (including residents of neighboring villages) is limited here. A farmer cultivating the land near the cave denied that this cave had been visited by any outsider.

**Table 2.** *Rousettus aegyptiacus* trapping data with the results of PCR screening on MARV.

| Roosting cave | Number of *R. aegyptiacus* / Number of MARV+ |  |  |
| --- | --- | --- | --- |
|  | Captured in the entrance of the cave | Captured on flight paths | Total |
| **Legon Tyo cave-related animals** | **13 / 1** | **6 / 1** | **19 / 2** |
| Juveniles | 4 / 1 | 2 / 0 | 6 / 1 |
| Adults | 9 / 0 | 4 / 1 | 13 / 1 |
| **Kokongoa cave-related animals** | **14 / 1** | **26 / 0** | **40 / 1** |
| Juveniles | 1 / 1 | 8 / 0 | 9 / 1 |
| Adults | 13 / 0 | 18 / 0 | 31 / 0 |
| **Animals for which the roosting cave was not found** | **0 / 0** | **7 / 0** | **7 / 0** |
| Juveniles | 0 / 0 | 4 / 0 | 4 / 0 |
| Adults | 0 / 0 | 3 / 0 | 3 / 0 |
| Total | 27 / 2 | 39 / 1 | 66 / 3 |

#### Kokongoa cave

(N 8.64561°, W 10.37876°) is located 48 km away from Temessadou M’bokét village on the northeastern slope of Mount Kokongoa (Tekoulo subprefecture). The cave has a wide tunnel inside with two large entrances: the upper and lower entrances are located at elevations of 642 and 605 meters above sea level, respectively. People from the surrounding villages visit this cave for ritual sacrifices at least once a year (usually in January) and store their ceremonial material directly at the lower exit of the cave. A visual inspection of the cave, as well as trapping with mist nets installed at both entrances, revealed the presence of a colony of ERB in the cave, as well as *Hipposideros ruber/caffer, Hipposideros beatus*, and *Hipposideros jonesii*. A total of 40 *R. aegyptiacus* were caught at the cave entrance and in the surrounding area (Table 2).

Seven ERB were also caught on flight paths in three different locations (Fig 1), but we did not find a roosting cave for these fruit bats. Additionally, we caught bats in 11 locations within a 20 km radius of Temessadou M’bokét village, including three caves (Fig 1), but other ERB colonies were not found in this area in either the caves or the flight paths.

**Fig 1.**
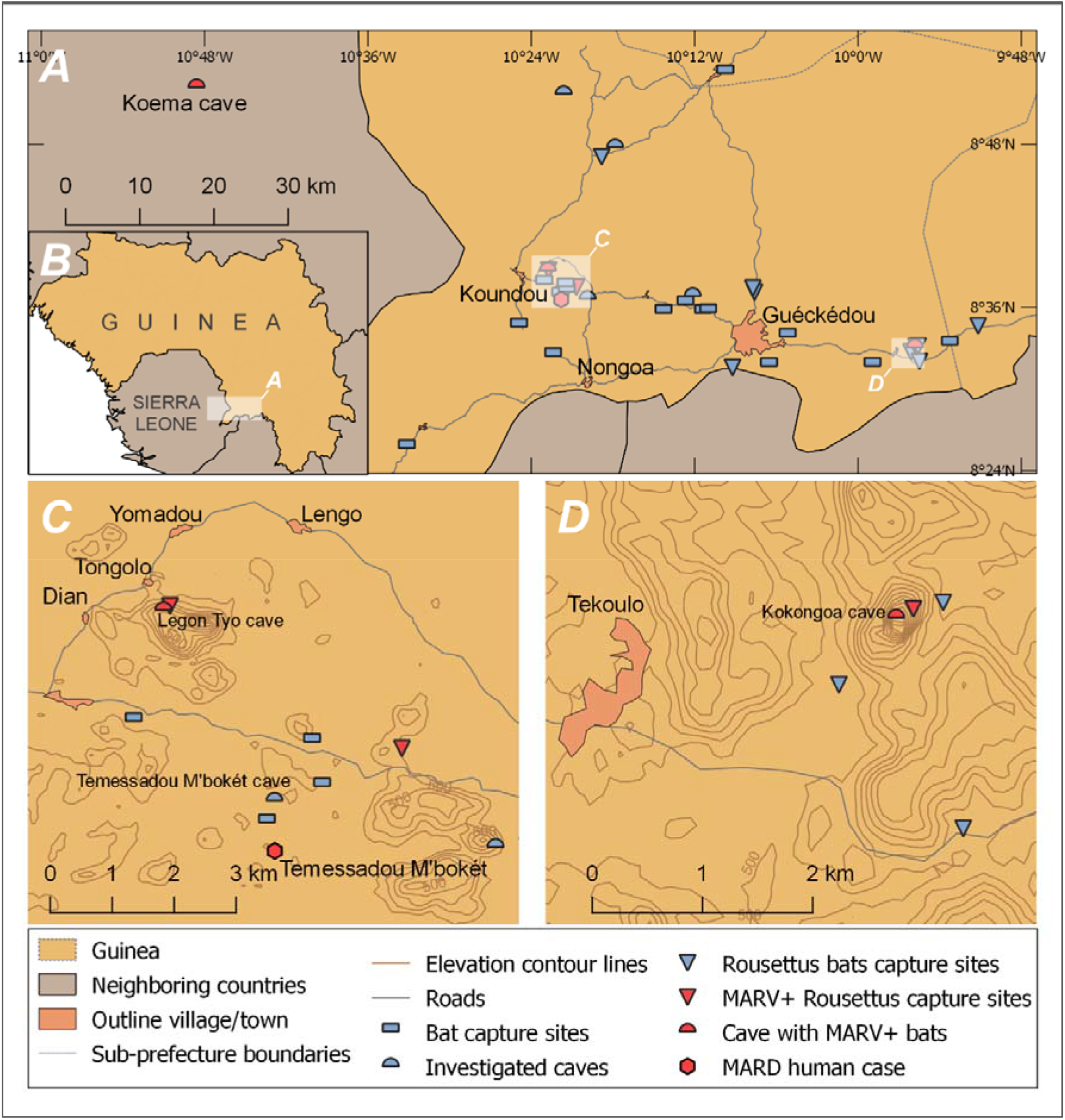
Sampling locations depicting the caves where PCR-positive MARV bats were caught. Map A shows the sampling locations in Guéckédou prefecture (Guinea) indicating the Koema cave in Sierra Leone where MARV-positive bats were found in a previous work [2]. Map B shows a general view of the sampling area and Guinea. Map C focuses on the area around Legon Tyo Cave and Temessadou M’bokét village. Map D zooms the area around the Kokongoa cave. Elevation contours on maps C and D are shown for altitudes over 440 m with an interval of 20 m.

### PCR detection of MARV in *Rousettus aegyptiacus* and Sanger sequencing

During the screening stage, homogenates of lymph nodes and pools of sections of internal organs from all trapped ERB were tested in the presence of MARV RNA. All samples with the exception of three were found to be negative. A total of three positive samples were obtained: two of these bats were trapped at the entrance or in the neighborhood of Legon Tyo cave, and one was trapped at the entrance to Kokongoa cave (Table 2). qRT□PCR showed very low viral loads in bat organs and tissues (Table 3). The lowest Ct value was in juvenile female ERB from Legon Tyo cave (bat id: 452), particularly in the spleen tissue and axillary lymph nodes (Ct values 27.1 and 28.1, respectively) (Table 3). We collected four bat flies, *Eucampsipoda africana*, from this animal and tested them separately. All four bat flies were PCR-negative for MARV. For the other two MARV-positive ERBs, the Ct values were both approximately 35. Oral swabs of all three PCR-positive ERB gave negative MARV results.

**Table 3.** Detailed data on MARV-positive *Rousettus aegyptiacus*.

| Location | Bat ID | Sex | Age status | RT-PCR Ct value | Sequencing data |
| --- | --- | --- | --- | --- | --- |
| Legon Tyo | 79 | male | Adult | Lymph node – 35.0<br>Lung, kidney, liver, spleen – negative | GenBank accession number: OP729425 |
| Legon Tyo | 452 | female | Juvenile | Spleen – 27.1<br>Liver – 32.2<br>Kidney – 32.1<br>Lymph node, axillary – 28.1<br>Lymph node, mesenteric – 31.8<br>Salivary glands – 30.3<br>Ectoparasites*, oral swab, brain, lung – negative | GenBank accession numbers: OP716850-OP716853 |
| Kokongoa cave | 387 | female | Juvenile | Liver+spleen – 35.2<br>Lung, kidney, lymph node – negative | No |

\* – four bat flies (*Eucampsipoda africana*) were collected and tested

We obtained sequences for fragments of the nucleoprotein (NP), viral protein 35 (VP35), viral protein 30 (VP30), and polymerase (L) genes with a total length of 3585 bp for the RNA isolated from the bat with ID 452 (Legon Tyo cave).

Phylogenetic analysis of the concatenated gene fragments showed that the identified isolate is most similar (p-distance = 0.012) to the sequence of MARV obtained in Sierra Leone from ERB (GenBank accession number MN258361, Fig 2).

**Fig 2.**
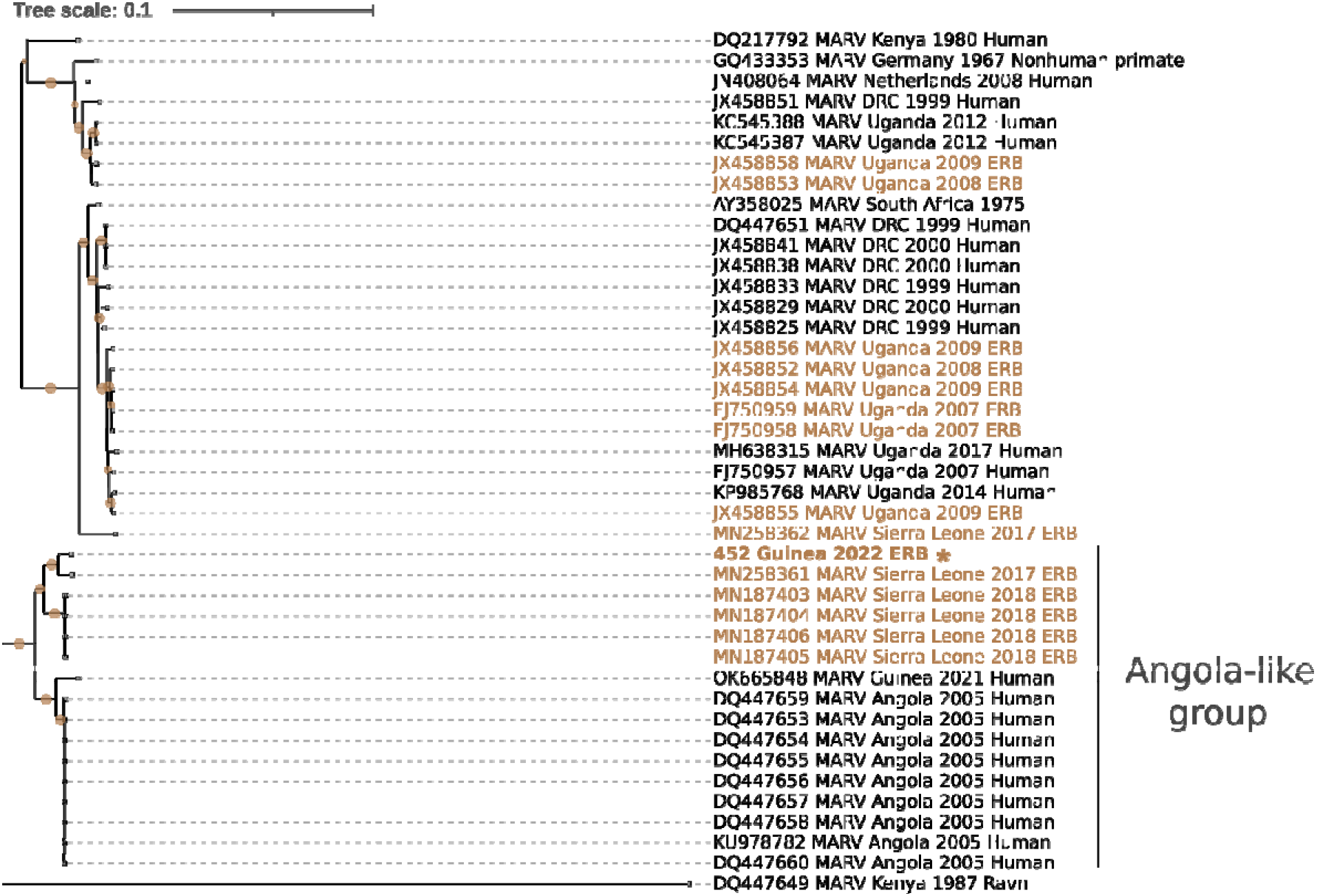
Maximum-likelihood phylogeny of 42 partial and concatenated Marburg virus nucleoprotein (NP, 916 bp), viral protein 35 (VP35, 300 bp), viral protein 30 (VP30, 1107 bp), and polymerase (L, 1262 bp) gene fragments with a total length of 3585 bp. The tree was constructed using the best-fit nucleotide substitution model (GTR+I+G). Sequences in brown represent those generated from ERB, and sequences in black represent those generated from human and nonhuman primate samples. The sequences from *R. aegyptiacus* ID 452 generated during this study are indicated with an asterisk and bold text. The scale bar indicates the mean number of nucleotide substitutions per site. The filled circles on branches indicate bootstrap values greater than 0.9.

For bat with ID 79 (Legon Tyo cave), we sequenced only the fragment of 92 bp of the NP gene. This short sequence allowed us to confirm that we did detect RNA of MARV and that the isolate differed from sample 452 (Fig 3). Surprisingly, although these two PCR-positive bats were supposed to belong to the same colony, the sequence of sample 79 did not belong to the Angola-like group and aligned most closely with MARVs of major marburgvirus lineages (clustered with isolates from Uganda and DRC) (Fig 3). The closest sequence was the virus previously found in ERB in Sierra Leone (GenBank accession number MN258362).

**Fig. 3.**
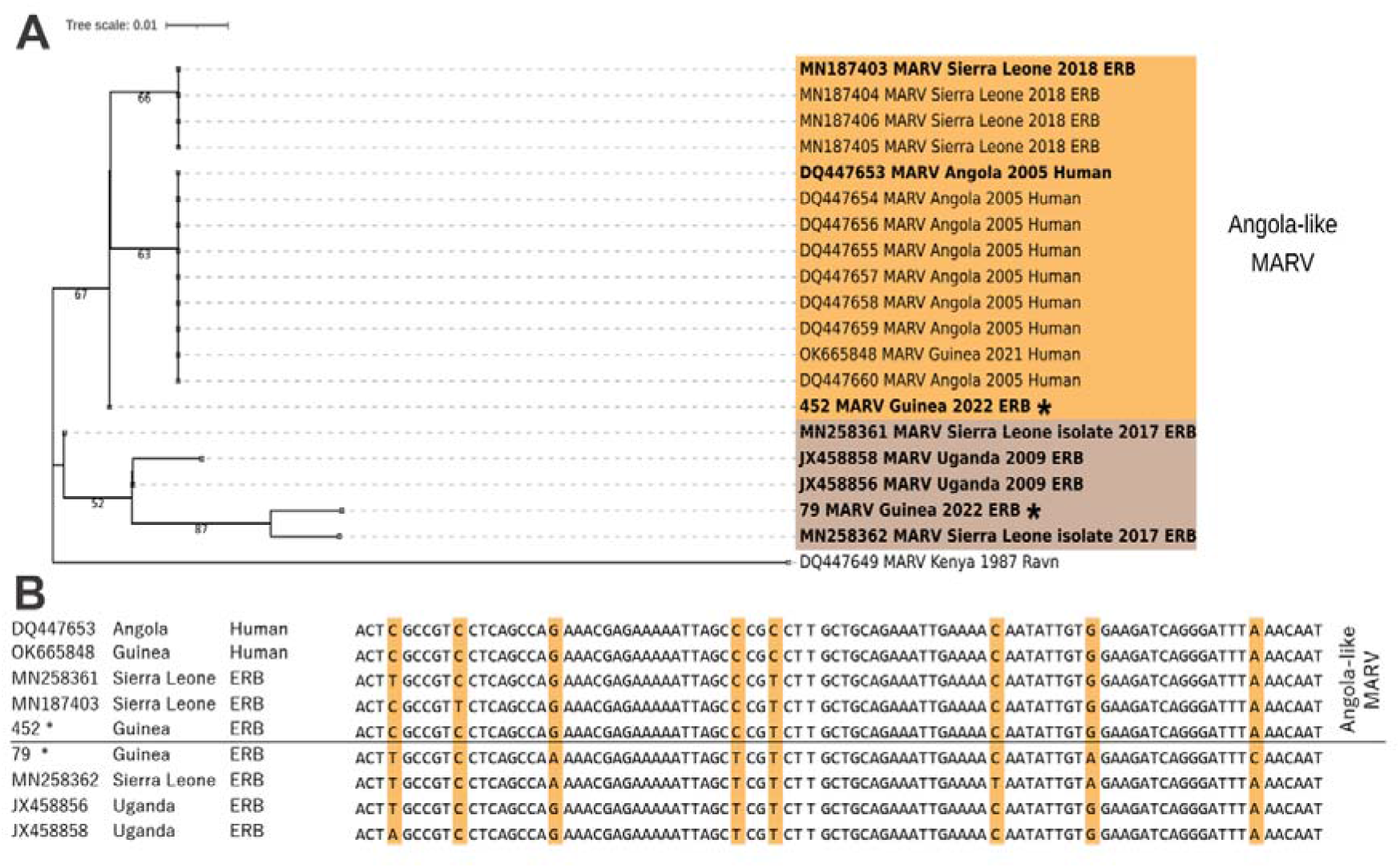
Phylogenetic tree and alignment comparing the relationship of the MARV sequence of sample ERB_79 to other marburgviruses. The sequences from Guinea generated during this study are indicated with asterisks. **A:** Maximum-likelihood tree (Tamura 3-parameter model with 1000 pseudoreplicates) of partial marburgvirus nucleoprotein gene fragments (92 bp). The scale bar indicates the mean number of nucleotide substitutions per site. The sequences with bold type were used in the alignment presented below. **B:** Aligned fragments (92 bp) of the MARV nucleoprotein gene with highlighted variable sites.

We were unable to obtain sequences from specimen 387 (Kokongoa cave).

## Discussion

In this work, we confirm the presence of MARV in ERB in Guinea. Moreover, we found an MARV-positive ERB in close proximity (4.5 km) to the village Temessadou M’bokét, where an MARD outbreak emerged in 2021. However, this patient could not have been infected while visiting this cave, as it is inaccessible to humans. We consider the two most likely MARV spillover scenarios in the 2021 outbreak. The first is direct contact with an infected ERB outside the cave while opportunistically hunting, catching, or finding a weakened animal in places of flight and/or feeding. This assumption is supported by the work of Saéz et al. [20], which showed that the local people living in Guéckédou prefecture eat bats, hunting them with guns, or opportunistically hunting with sticks or machetes and even catching them by hand. The second assumption is indirect spillover through consumption of infected fruits or through contact with an ill primate that had become ill after eating contaminated fruits. ERB do not typically consume fruit completely; rather, they chew the fruit to extract the juice and then discard the pulp [24,25]. Additionally, they may urinate on fruit and test bite fruits or drop fruit they have been actively eating [6,24]. This feeding behavior, combined with the ability of infected ERB to shed the virus in urine and saliva [26–29], means that MARV can be spread through fruits that ERB have previously eaten. Amman et al. [24] confirmed this plausible route of MARV transmission with an experiment and showed that MARV is stable for at least six hours on fruits. Based on the above, we can consider that such a spillover scenario is also very likely for the 2021 MARD outbreak. It is possible that there was another ERB colony in the vicinity of Temessadou M’bokét village that roosted in a cave that we could not find. However, our survey of local residents showed that they do not know of other caves in the area. Until other caves are discovered, the ERB colony in Legon Tyo cave is the closest source of the MARV to Temessadou M’bokét village (4.5 km). Assuming that the patient from the village got the virus from the ERB of the Legon Tyo colony, then the 2021 MARD outbreak emerged after contact with the virus outside the cave.

The detection of MARV-positive fruit bats in the Kokongoa cave, far from the Legon Tyo cave (50 km) and Temessadou M’bokét village (48 km), indicates that MARV is most likely to be more widespread in Guinea and that there are likely other caves in which the ERB roost was infected. Perhaps there have been other outbreaks of MARD in Guinea before, but they just were not diagnosed. Therefore, we consider it important to study ERB colonies in other prefectures of Guinea (Nzérékoré, Beila, Macenta, Youmu, Lola, Madina Oula, Mamou, Kindia, etc.). Moreover, similar research should also be carried out in other countries of West Africa (in Liberia, Côte d’Ivoire, Mali, Burkina Faso, etc.). The 2022 Ghana MARD outbreak showed that MARV is more widespread in West Africa than previously known.

We caught three MARV-positive ERB, and two of them were juveniles. Due to the lack of samples, we could not claim an age bias. However, Amman et al. [2] showed a significant age bias among MARV-positive bats in neighboring Sierra Leone: all 11 PCR-positive bats were juveniles. Furthermore, it was confirmed previously that juvenile ERB are more frequently infected by MARV than adult animals [12,14]. Therefore, we can expect a similar age bias in Guinea, with a predominance of juveniles among MARV-positive ERB. Thus, to prevent possible new outbreaks of MARD, it is important to know not only the location of the caves but also the seasonality of ERB breeding [12]. In sub-Saharan Africa, the ERB has biannual birthing seasons [6,30]. We assume that in Guinea, as well as in Uganda, the reproductive chronology of ERB is seasonal bimodal polyestry without postpartum estrus [31,32]. Females have two litters per year and a gestation of 106 days [6]. Females are in reproductive synchrony; therefore, young are born in two distinct seasons of parturition. Based on observations in Liberia [33–35] and Sierra Leone [2], fruit bats in Guinea have two parturition seasons, with peaks in December and June. After three months, the puppies fly out of the cave on their own for the first time and gradually begin to explore the area around the cave [36]. At this time, juveniles pose the greatest danger, as they significantly increase the likelihood of a spillover event. Amman et al. [12] showed that juvenile ERB approximately 4.5–7.5 months old are most likely to be infected with MARV. According to our assumptions in Guinea, this period of greatest risk falls on April-July and October-January. This correlates with the time of the outbreak in 2021: the first MARD symptoms in the patient appeared on July 25 [1]. This is also consistent with data from Sierra Leone, where MARV-positive juveniles were caught in September-October and December [2].

In PCR screening, we obtained low concentrations of MARV in tissues, with Ct values above 27 (Table 2). Similar Ct values (>28) were obtained in other studies [2,5]. This result highlights the difficulty of detecting MARV in field studies and shows the high requirements for sample storage, transport and processing.

Sequencing of two virus amplicons obtained from ERB caught in the Legon Tyo cave and on flyways located near the cave and Temessadou M’bokét village (4.5 km and 2.6 km, respectively) showed that different genetic variants of MARV circulate here: Angola-like MARV and MARV of major marburgvirus lineages (Uganda, DRC genetic variant). Wholegenome sequences of these two genetic variants differ from each other by approximately 7% [2]. Similar data on genetic diversity were obtained for the MARV found in EBR in Sierra Leone, where two different genetic variants of MARV were found in the ERB colony of Kasewe Cave: Angola-like MARV and MARV phylogenetically close to isolates from Gabon, DRC, and Uganda [2].

MARV sequences from ERB of the Legon Tyo colony are not identical to the MARV isolate from the Temessadou M’bokét patient. The MARD patient isolate is close to sequences from the 2005 Angola outbreak with 98% identity and differs by 3.5% from specimen ERB-452 (Legon Tyo ERB colony). A possible explanation for this difference is the small number of sequenced PCR-positive ERB. Another hypothesis is that there are at least two genetic lines within the Angola-like MARV: the first is more adapted to ERB, and the second is adapted to humans.

To summarize, the information obtained is sufficient to take measures to prevent possible new spillovers: the exact location of the cave where MARV-positive ERB roost was determined. Furthermore, it is likely that the MARV spillover event can occur not only when visiting the cave but also within the foraging radius of the ERB. We would recommend to local authorities to inform the inhabitants of the surrounding villages that fruit bats live in their area and are infected with a very dangerous virus; therefore, they should not be caught and eaten. In addition, to prevent possible spillover through contaminated fruit, it is necessary to wash any fruit before consumption. We would also recommend avoiding contact with sick and dead monkeys, which can also be a source of MARV.

## Conclusion

In this work, we showed the presence of PCR-positive MARV fruit bats in Guinea. Sequencing has shown that at least two different MARV genetic variants circulate in ERB in Guinea: an Angola-like MARV and MARV of major marburgvirus lineages. We have found and described two caves in Guéckédou Prefecture where MARV-positive ERB roost. Our findings suggest that MARV is more widely distributed in Guinea than in a single location in Guéckédou prefecture, and screening is needed in other parts of the country. These results provide the basis for preventive measures against new outbreaks in Guinea.

